# A practical method for efficient and optimal production of selenomethionine-labeled recombinant protein complexes in the insect cells

**DOI:** 10.1101/491548

**Authors:** Sabine Wenzel, Tsuyoshi Imasaki, Yuichiro Takagi

## Abstract

The use of Selenomethionine (SeMet) incorporated protein crystals for single or multiwavelength anomalous diffraction (SAD or MAD) to facilitate phasing has become almost synonymous with modern X-ray crystallography. The anomalous signals from SeMets can be used for phasing as well as sequence markers for subsequent model building. The production of large quantities of SeMet incorporated recombinant proteins is relatively straightforward when expressed in *E. coli*. In contrast, production of SeMet substituted recombinant proteins expressed in the insect cells is not as robust due to the toxicity of SeMet in eukaryotic systems. Previous protocols for SeMet-incorporation in the insect cells are laborious, and more suited for secreted proteins. In addition, these protocols have generally not addressed the SeMet toxicity issue, and typically result in low recovery of the labeled proteins. Here we report that SeMet toxicity can be circumvented by fully infecting insect cells with baculovirus. Quantitatively controlling infection levels using our Titer Estimation of Quality Control (TEQC) method allows for incorporation of substantial amounts of SeMet, resulting in an efficient and optimal production of labeled recombinant protein complexes. With the method described here, we were able to consistently reach incorporation levels of about 75% and protein yield of 60-90% compared to native protein expression.

## Introduction

Single or multi-wavelength anomalous diffraction (SAD or MAD) phasing by utilizing Seleno-methionine (SeMet) labeled proteins or protein complexes is almost synonymous with modern X-ray crystallography ^1,2^. This method has drastically reduced the need for preparation of heavy atom derivatives, which is one of the most time-consuming steps in X-ray crystallography. The anomalous signal from SeMet can be used for phasing as well as for identifying methionine positions within the electron density map, thereby aiding model building in low resolution maps ^3,4^

Production of SeMet-labeled proteins in *E. coli* for X-ray crystallography is well established and robust ^1^. Protocols exist for yeast *Saccharomyces cerevisiae* ^3^, yeast *Pichia pastoris* ^5^, the insect cells ^6,7^ as well as mammalian cells ^8^. While SeMet incorporation routinely achieves 100% for proteins expressed in *E. coli*, expression in eukaryotic cells is much more variable, with SeMet incorporation ranging from 50 % to 90 %, depending on the host cells and proteins being produced ^5,7–10^.

The baculovirus expression system (BEVS) is an excellent tool for expressing recombinant proteins in insect cells. It is considered an attractive option for proteins as well as protein complexes and has been widely used for structural and functional studies ^11–13^. For production of SeMet-labelled proteins using the BEVS in the insect cells, the protocol by Bellizzi et al. has been widely cited ^6^. However, as pointed out by Cronin et al. ^7^, this protocol is suited for secreted proteins such as glycosylated lysosomal hydrolase ^6^, and CD40 ^14^, and is not necessarily applicable for intracellular expression. Moreover, this protocol requires several medium-exchanges and dialyzed FBS added to the methionine-free medium in order to maintain cell viability during methionine depletion. Since higher labor demand (more personnel cost) as well as requirement of additional reagents such as FBS will add to the overall cost of operation, this protocol could potentially become costly. To overcome some of these limitations, a protocol designed for secreted as well as intracellularly expressed proteins was published ^7^. However, this protocol still retains the cumbersome medium exchange in the midst of the operation, and yields were drastically reduced: 18%-45% compared to the native protein. Low yield of SeMet-labeled proteins is a serious problem as it limits crystallization conditions for screening, optimization, and production of crystals, and low incorporation rates reduce the available anomalous signal. Moreover, additional preparations are required to achieve desired protein yield leading to increased costs as well.

In our work on the yeast Mediator Head module, which is an essential subcomplex of the Mediator complex, and a key component of transcription regulation in eukaryotes ^15^, we encountered a similar problem. The Mediator Head Module is composed of seven subunits with a total molecular mass of 223 kDa. We generated the recombinant complex in the insect cells using the MultiBac baculovirus expression vector system for structural and functional studies ^4,16^. We wanted to use the SeMet-labeled Mediator Head module for phasing as well as for model building purposes. Using a protocol similar to that from Cronin et al. ^7^ or the protocol provided by the manufacturer (Expression Systems, Inc), we achieved only 10% or less recovery of the Mediator Head module compared to the native expression level. A substantial reduction of the SeMet-labeled protein yield could likely be attributed to SeMet toxicity. Although this toxicity is wildly recognized in the field, there is no detailed experimental data available as to toxic dosages of SeMet to the insect cells. Chen and Bahl (1991) reported that 100 μg/ml of SeMet was toxic but they did not present dose-dependent toxicity data, or experimental details ^17^. Further, there has been no investigation undertaken of potential ways to bypass or reduce the detrimental effects of SeMet on the insect cells, as it had been published for yeast S. *cerevisiae* ^10, 18^.

Considering the limitations and intricacy of the current SeMet-labeling methods in the insect cells ^6,7,17^, we decided to analyze critical aspects of SeMet incorporation in the insect cells in order to improve quality and quantity of SeMet-labeled proteins. We report here that SeMet is indeed toxic to the insect cells, but such toxicity can be circumvented by high baculovirus infection levels. We developed a simple, quantitative, and easy-to-use protocol for the generation of SeMet-labeled proteins or protein complexes in the insect cells.

## Results

### Sf9 and Hi5 insect cells are resilient to methionine depletion

We started off investigating the effect of methionine depletion on cell viability in Sf9 and Hi5 cells. We wanted to test if supplementing dialyzed FBS for cell maintenance under met-depleted conditions ^6, 17^ is necessary. Both cell lines were maintained in serum-free ESF921 medium (Expression Systems, Inc). Cells, Hi5 or SF9, were spun down and resuspended in either the Met-containing ESF921 medium (control) or Met-free ESF921 medium in a 50 ml culture with a cell density of 0.5 x 10^6^ (Hi5) or 0.6 x 10^6^ (Sf9) cells/ml. Cell viability (% of live cells) and cell density of each culture was monitored every 24 hours for a total of 4 days (Figure 1). Although growth of Hi5 cells in the methionine-free medium began to be compromised on day 3, Sf9 cells remained the same as the control culture during the 4-day incubation period. More importantly, there is essentially no difference in cell viability with or without methionine in both cell lines (Figure 1A, 1C), suggesting strongly that both insect cell lines are resilient to methionine depletion for an extended period of time: Sf9 for at least 96 hours, and Hi5 for 48 hours. Therefore, we concluded that (i) addition of dialyzed FBS during methionine depletion is unnecessary, and that (ii) methionine depletion can be sustained for at least 48 hours for Hi5 cells, and 96 hours for Sf9 cells.

**Figure 1:**
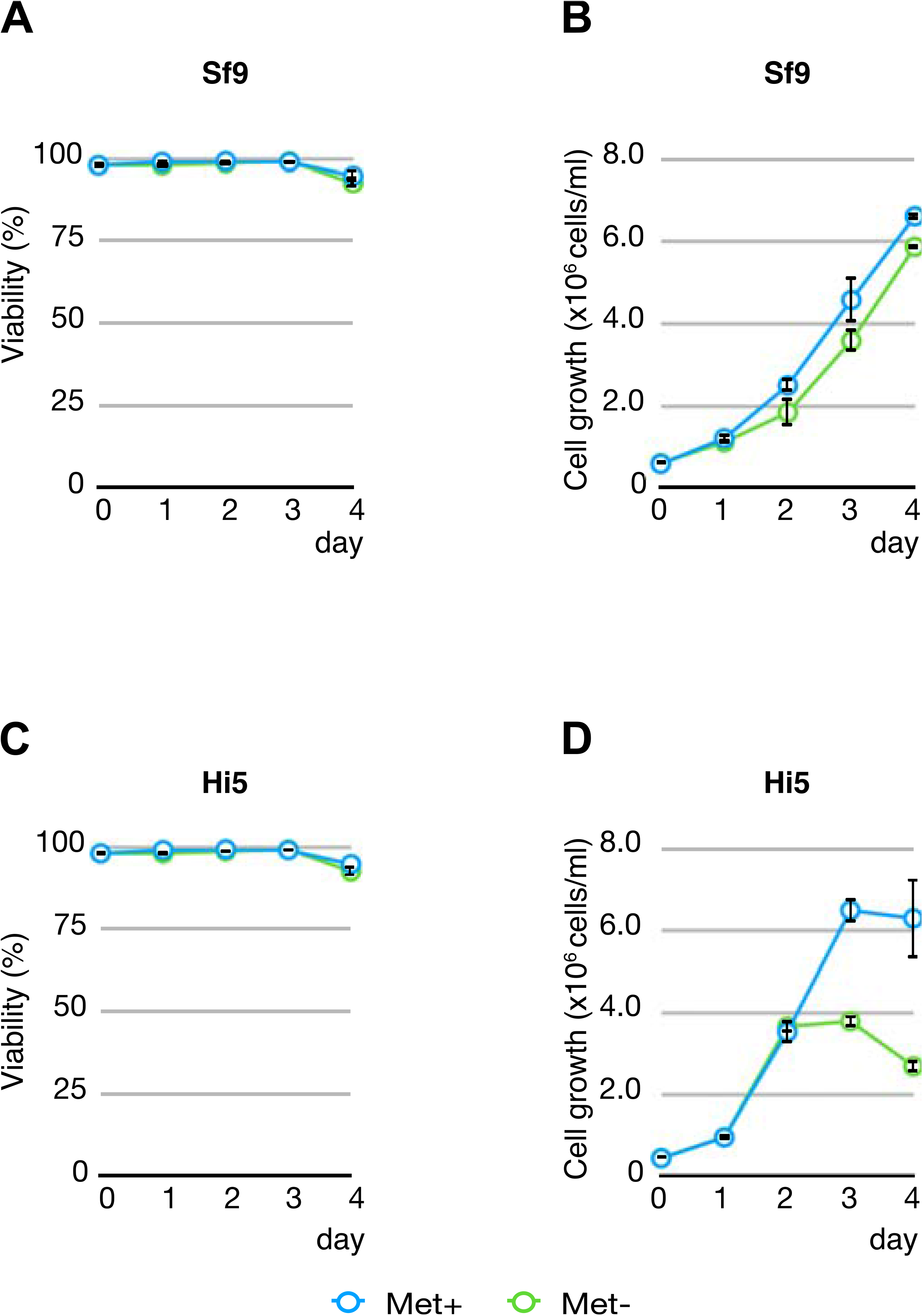
Effect of methionine depletion on Sf9 and Hi5 insect cells. Hi5 and Sf9 cells were grown in either Met-free or Met-containing medium as control. Cell viability (% of live cells) and cell density of each culture was measured every 24 hours for a total of 4 days. (A) Cell viability of Sf9 cells with or without Met (B) Cell density of Sf9 cells with or without Met (C) Cell viability of Hi5 cells with or without Met (D) Cell density of Hi5 cells with or without Met. Measurements were performed in triplicates and averaged. Cyan: Met-containing medium; green: Met-free medium

### SeMet is toxic to the insect cells

Next, we examined SeMet toxicity in the insect cells in the presence or absence of Met by monitoring cell viability as well as overall cell growth. Sf9 or Hi5 cells were passaged with fresh ESF921 medium (Met+, control) or Met-free ESF921 medium (Expression System, Inc) to a cell density of 0.5 x 10^6^ cells/ml in a 50 ml culture. Various amounts of SeMet, ranging from 0 mg/L to 200 mg/L, were added to both sets of cultures (Met+ or Met-). Cell viability (% of live cells) and cell density (total cell density) of each culture was monitored every 24 hours for 4 days (Figure 2). In the presence of Met, viability and growth of Sf9 cells were only slightly affected by SeMet at a concentration of 20-80 mg/L; at 120-200 mg/L, cell viability began to decrease (Figure 2A, 2B). Hi5 cells are more susceptible to the presence of SeMet, and their viability is dependent on the SeMet concentration in the medium (Figure 2E, 2F). In the Met-free medium, the toxic effect of SeMet appears to manifest itself more prominently such that the cell viability drops to 40% in Sf9 cells (Figure 2C) or 18% in Hi5 cells (Figure 2G) at 80 mg/L SeMet. At 80 mg/L or higher, cell viability of both cells declined to ~20% (Figure 2C, 2G), and cell growth of both cell lines was severely compromised (Figure 2D, 2H). These data clearly suggest that SeMet has detrimental effects on cell viability and cell growth *per se*, and impacts cell culture growth significantly at concentrations of 80 mg/L or higher in Met-free medium. Based on the data presented here, the amounts of SeMet used in several publications ^6,7,14,17,19,20^ clearly fall in the toxic range.

**Figure 2:**
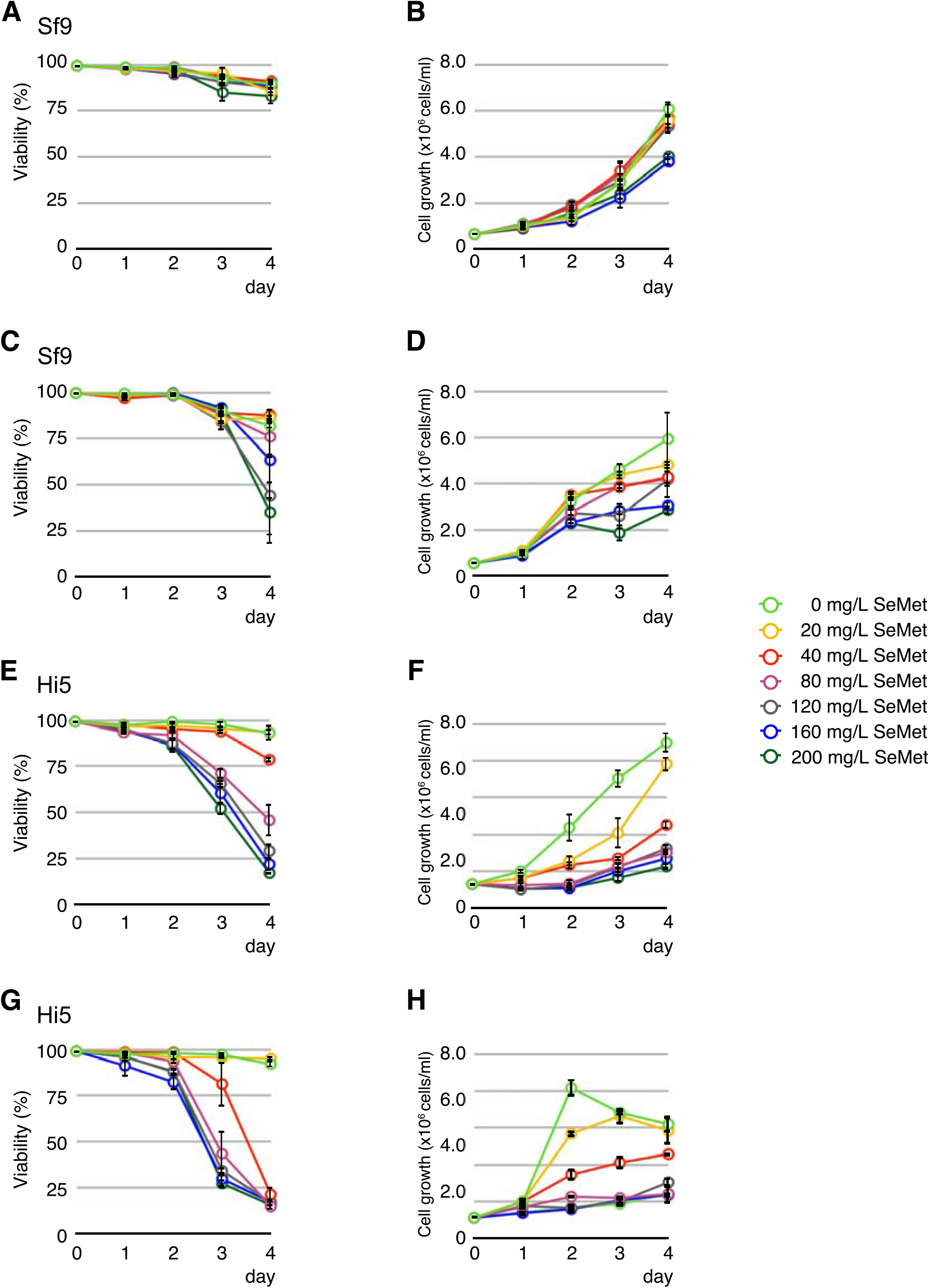
Effect of SeMet on Sf9 or Hi5 insect cells cultured in Met-containing medium, or Met-free medium. Hi5 or SF9 cells were seeded in Met-containing or Met-free medium followed by addition of SeMet to the final concentration of 0 (control), 20, 40, 80, 120, 160, and 200 mg/L, respectively. Cell viability (% of live cells) and cell density of each culture was measured every 24 hours for a 4-day span. (A) Cell viability of Sf9 cells in Met-containing medium (B) Cell growth of Sf9 cells in a Met-containing medium (C) Cell viability of Sf9 cells in Met-free medium (D) Cell growth of Sf9 cells in Met-free medium (E) Cell viability of Hi5 cells in a Met-containing medium (F) Cell growth of Hi5 cells in a Met-containing medium (G) Cell viability of Hi5 cells in Met-free medium (H) Cell growth of Hi5 cells in Met-free medium. Final concentration of SeMet: green: 0 mg/L; yellow: 20 mg/L; red: 40 mg/L; magenta: 80 mg/L; gray: 120 mg/L; blue: 160 mg/L; dark green: 200 mg/L

### SeMet toxicity can be circumvented by baculovirus infection

Having established that the insect cells are resilient to Met depletion for an extended period of time (Figure 1), and that SeMet is indeed toxic to the insect cells at concentrations at 80 mg/L or higher under Met depleted conditions (Figure 2), we investigated the relationship between SeMet incorporation, and the yield of the SeMet-labeled Mediator Head module. For this purpose, we set up cultures of Hi5 cells infected with the Mediator Head module virus at an estimated multiplicity of infection (eMOI) of 3.7, which gave consistently high yield in a native expression as we reported previously ^21^, and titrated increasing amounts of SeMet into the cultures (Figure 3). In this experiment, Hi5 cells were chosen because the Mediator Head module expressed significantly better in Hi5 cells than Sf9 cells ^21^. The overall experimental scheme is described in Figure 3A. For ease of implementation, our labeling protocol was divided into 24 hours (one day) increments. On Day 0, Hi5 cells grown in log phase were spun down and resuspended in Met-free medium. Cultures were incubated for 24 hours to deplete Met, and after adjusting for cell density, the baculovirus expressing the Mediator Head module was added to each culture (Day 1) with an eMOI=3.7 ^21^. On Day 2, 0 (control) to 200 mg/L SeMet was added to each culture and 100mg/L Met was added to the native control culture. Cell density and viability were monitored every 24 hours over a 4-day period (Day 1-5). To our surprise, we saw no significant SeMet toxicity as assessed by cell density and viability over the measured time period (Figure 3B, 3C): Cell viability at 200 mg/L SeMet (52%) was similar to that of the native control with 100 mg/L Met (53%) added (Figure 3B). Our previous work showed that Hi5 cells were fully infected with an eMOI=3.7 ^21^. Thus, we hypothesize that high baculovirus infection levels could circumvent the SeMet toxicity in the insect cells.

**Figure 3:**
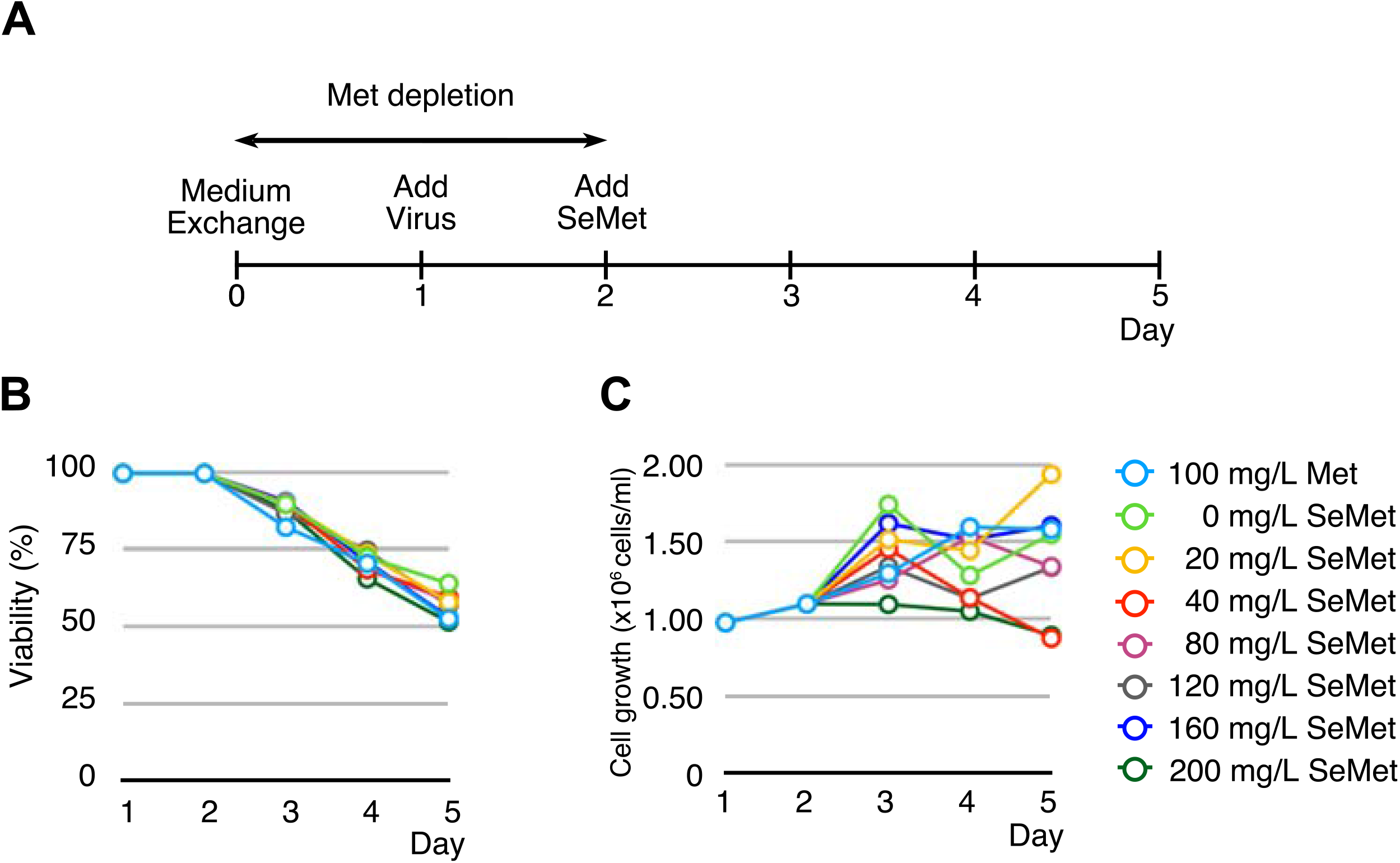
Effect of baculovirus infection on SeMet toxicity. (A) Overall experimental scheme. On Day 0, culture medium was exchanged from Met-containing to Met-free to deplete intracellular methionine. On Day 1, the baculovirus expressing the Mediator Head module with an eMOI=3.7 was added to the culture. On Day 2, SeMet was added to each culture to the final concentration of 0 (control), 20, 40, 80, 120, 160, and 200 mg/L, respectively. As for the control, native expression was set up such that Met with a final concentration of 100 mg/L was added. Cell viability (% of live cells) (B), and cell density (C) of each culture was measured every 24 hours for a 4-day period (Day 1-5). Cyan: 100 mg/L of Met, for SeMet, green: 0 mg/L; yellow: 20 mg/L; red: 40 mg/L; magenta: 80 mg/L; gray: 120 mg/L; blue: 160 mg/L; dark green: 200 mg/L

### Cell viability and recovery of the SeMet-labeled Mediator Head module in the presence of a toxic amount of SeMet depends on infectivity (or eMOI)

If a high baculovirus infection level could evade the SeMet toxicity, thereby maintaining cell viability, it should lead to higher recovery of SeMet-labeled proteins or protein complexes. Therefore, we further hypothesized that optimal baculovirus infection is as critical for optimal SeMet-labeled protein yield as it is for cell viability in terms of evading SeMet toxicity: optimal baculovirus infection leads to optimal SeMet-labeled protein yield.

To test this hypothesis, we set up a virus titration such that the infectivity of the insect cells at the time of SeMet addition varied from low (22%) to high (~100% infected cells) in order to see if there is a correlation among infectivity, cell viability, and SeMet-labeled protein yield in the presence of a toxic amount of SeMet. We previously developed the Titer Estimation of Quality Control (TEQC) method, which enables us to quantitatively control virus infection levels ^21^. Using our TEQC method, the titer value of our virus stock was estimated and cultures with a total of five different estimated initial infectivities (infectivity 24 hours after addition of virus: el_24_), ranging from 22%, 39%, 63%, 86% to 98%, were set up. These initial infectivities correspond to estimates of multiplicity of infection (eMOIs) of 0.25 (22%), 0.5 (39%), 1.0 (63%), 2.0 (86%), and 4.0 (98%), respectively. It should be noted that the mathematical relationship between an initial infectivity (I_24_) and MOI is non-linear: I_24_=1-e^-MOI 21^. The corresponding eMOI, instead of infectivities, were used for the experiments in Figure 4, since this experimental condition is more convenient to set up. For instance, an eMOI=2 implies twice as much virus volume was added than for eMOI=1. As for controls, native expressions were set up in parallel using identical eMOI conditions. As shown previously, Hi5 cells are better suited for expression of the Mediator Head module ^21^, and thus, Hi5 cells were used for this experiment. Following the same experimental scheme (Figure 3A), after Met depletion and adjustment of cell density the baculovirus was added to each culture with eMOI from 0.25 to 4.0 (Day 1). On Day 2 (24 hours after virus addition), SeMet was added at a final concentration of 160 mg/L, which was shown to be highly toxic in the absence of infection (Figure 2G), or Met was added at a final concentration of 100 mg/L as a native expression. Cell viability of each culture was monitored every 24 hours for a 4-day period (Day 1-5) (native: Figure 4A; SeMet: Figure 4B). On Day 5, cells were harvested, and the Mediator Head module was purified and quantified. Comparison in cell viability between a native vs. SeMet containing expressions on Day 5 is displayed in Figure 4C.

**Figure 4:**
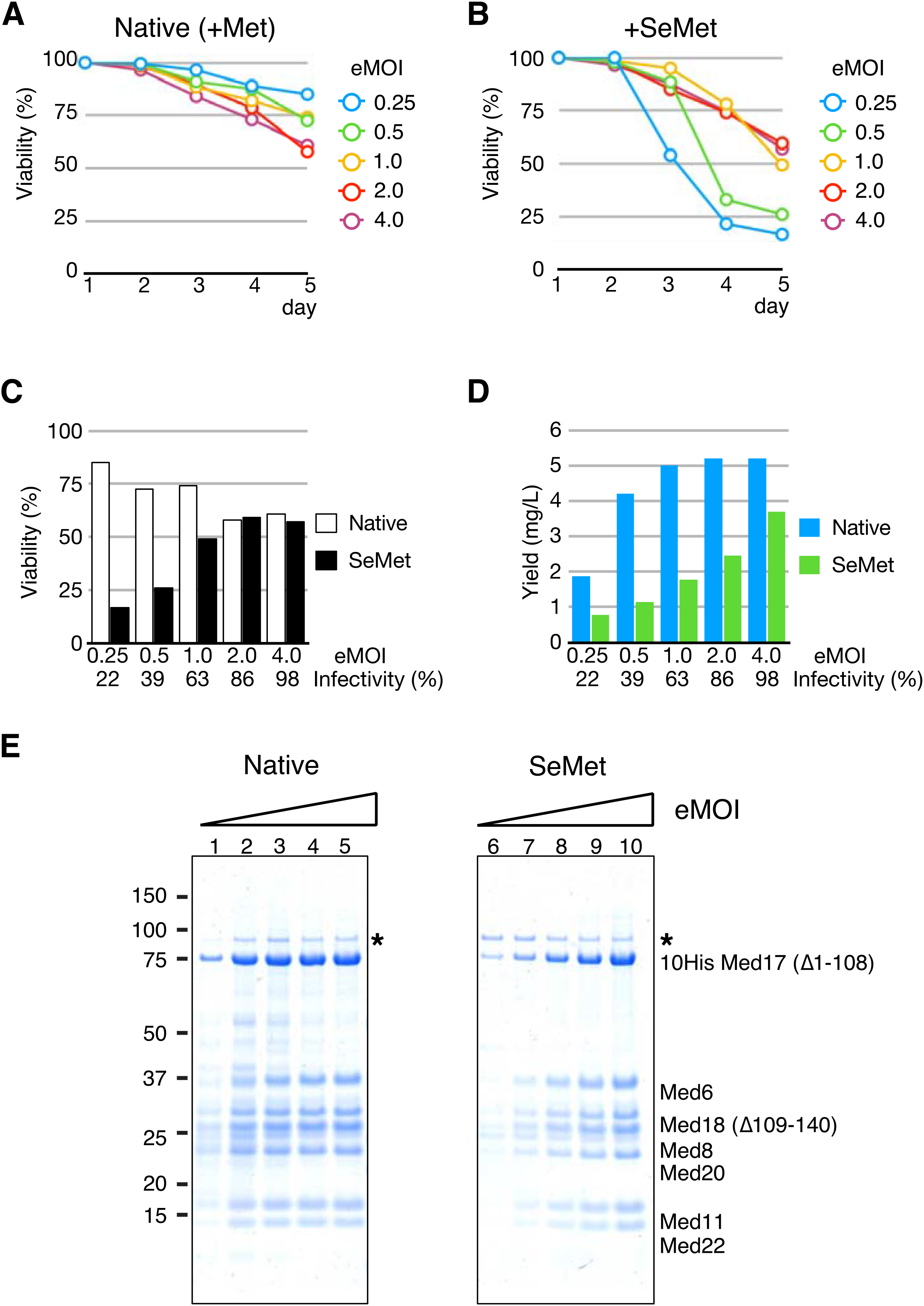
Cell viability and recovery of the SeMet-labeled Mediator Head module in the presence of SeMet depends on infectivity (or MOI) (A)-(B) Cell viability as a function of estimated multiplicity of infection (eMOI), or initial infectivity. Following the experimental scheme described in Figure 3A, after Met depletion process, the recombinant baculovirus expressing the Mediator Head module at eMOI ranging from 0.25 (cyan), 0.5 (green), 1.0 (yellow), 2.0 (red) and 4.0 (purple) was added on Day 1 followed by addition of Met at a final concentration of 100 mg/L as native expression control, or SeMet at a final concentration of 160 mg/L on Day 2. Cell viability at each condition was monitored during a 4-day period (Day 1-5), and cell viability (%) at each condition was plotted. (A) Native expression (B) SeMet labeling expression (C) Comparison of cell viability between native vs. SeMet containing expressions at Day 5 is displayed. White box: Native expression, Black box: SeMet labeling expression. (D) Recovery of the Mediator Head module in native or SeMet labeling expression condition as a function of estimated multiplicity of infection (eMOI), or initial infectivity. Cells were harvested on Day 5. The Mediator Head module was affinity-purified and quantified. The complex yield from each culture was measured and the data were plotted as a function of eMOI or initial infectivity. Yield was displayed as amount of the protein complex (mg) obtained from 1 L culture. Initial infectivities (%) correspond to eMOI were displayed below eMOI values. Cyan box: Native expression, green box: SeMet labeling expression (E) SDS-PAGE of the Mediator Head module obtained from the native expression on the left and SeMet-labeling expression on the right. (*): Contaminant from the insect cells

In the native expressions, cell viability went down to ~60% at eMOI greater than 2.0, which corresponded to the infectivity >86% on Day 5, while at low eMOIs (0.25, 0.5, or 1.0) or infectivity = 22%, 39%, or 63%, cell viability was in a range of 72-85% (Figure 4A, 4C). In the presence of SeMet (160 mg/L), cell viability was severely compromised at low eMOIs (0.25, or 0.5) or infectivity < 39% (Figure 4B, 4C). We speculate that this is due to a large percentage of cells being uninfected at the time of SeMet addition, and thus, far more prone to SeMet toxicity than infected cells. In contrast, at the higher eMOI range (2 or greater, or infectivity > 86%), cell viability went down to 57 % (Figure 4B, 4C), which was almost the same cell viability of the culture infected with eMOI=4.0 (infectivity 98% or higher) of the native expression (61%) (Figure 4C), indicating that cell viability at higher eMOI was not significantly affected under SeMet-labeling conditions: fully infected insect cells are far more resilient to SeMet toxicity than the uninfected.

Next, we looked into virus infectivity (or eMOI) and protein complex recovery. Protein yields were compared with those from native expression (Figure 4D). As for the native expression, protein complex yield appeared to reach a plateau at eMOI=1.0 or higher (Figure 4D, 4E). In contrast, in the presence of SeMet, highly infected cells (eMOI = 4) produced more protein complex than lower eMOI (Figure 4D, 4E). For example, the condition of eMOI=4 yielded 71 % of the protein complex compared to that of the native expression, whereas only 27 % of the native set up was recovered from cells infected with a low eMOI of 0.5. There is no significant difference in cell viability between cultures set up with eMOI =2.0, and 4.0. However, the protein complex yield recovered from the culture with eMOI=4.0 was ~34% better than that eMOI=2.0.

Taken all together, the data strongly suggests that a recovery of SeMet-labeled protein complex correlates with the infectivity (or eMOI) (Figure 4C, 4D): generating a condition of full infection (eMOI ≥ 4.0) at the time of SeMet addition should lead to optimal recovery of SeMet-labeled protein complexes.

### Optimal SeMet concentrations for SeMet labeling of the Mediator Head module expressed in Hi5 or Sf9 insect cells

After establishing the optimal conditions for evading SeMet toxicity without compromising recombinant protein recovery-adding the baculovirus with eMOI ≥ 4.0 - we investigated whether the higher tolerance for SeMet affects its incorporation into the recombinant protein complex. Ultimately, it is desirable to obtain high yield as well as a high SeMet incorporation. To study the effects of various SeMet concentrations on their incorporation into the Mediator Head module, Hi5 insect cells were cultured in Met-free medium followed by addition of the baculovirus expressing the Mediator Head module with an eMOI of 4 in order to fully infect cells. SeMet was added in concentrations ranging from 20 to 200 mg/L. As a control, a native expression test was set up in parallel with Met added at a final concentration of 100 mg/L. For each condition, the protein complex was purified and the yield was determined. The percentage of SeMet incorporation was measured by amino acid analysis (AAA). Consistent with the previously reported results ^7^, high concentration of SeMet resulted in higher incorporation of SeMet (Figure 5A). The SeMet incorporation gradually increased and peaked at 160 mg/L with 75% incorporation (Figure 5A). Consistent with the data in Figure 4C, the recovery of the protein complex was comparable to native levels up to a SeMet concentration of 160 mg/L (Figure 5B). The peak recovery of the complex at 160 mg/L SeMet is about 75% of that of native protein yield. No increase in incorporation level was observed at 200 mg/L SeMet (Figure 5A), but Mediator Head Module recovery was severely compromised at this concentration (Figure 5B). We concluded that for Hi5 cells, the maximum concentration of SeMet is 160 mg/L without overly compromising the recovery of the protein complex and achieve 75% SeMet incorporation.

**Figure 5:**
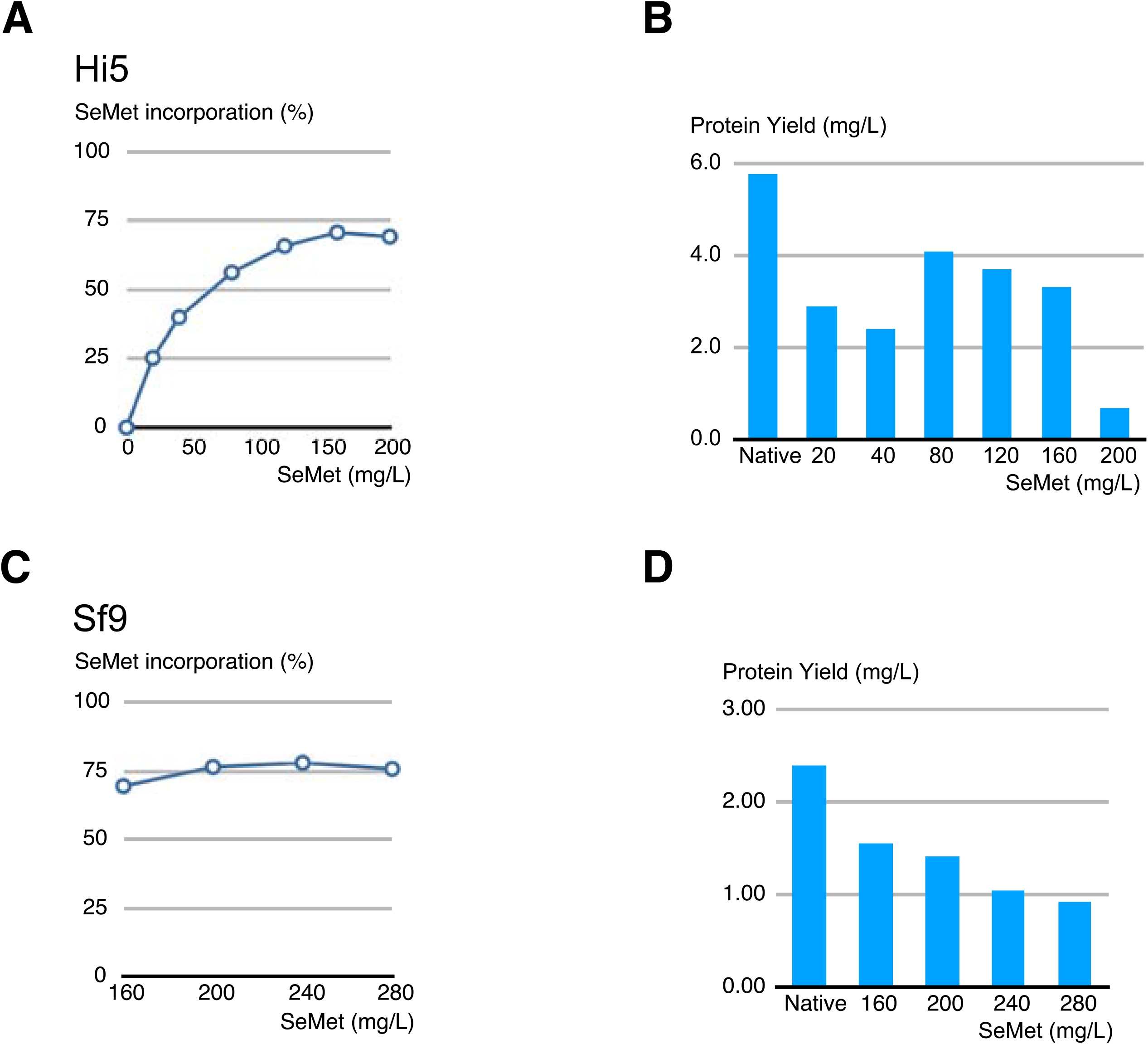
Effect of SeMet concentration on SeMet incorporation rate as well as on recombinant Mediator Head module recovery. Following the experimental scheme described in Figure 3A, Hi5 or Sf 9 cells (200 ml in 1-L shake flask) underwent the Met depletion process followed by the addition of the recombinant baculovirus expressing the Mediator Head module at eMOI of 4.0 on Day 1. On Day 2, SeMet was added to various final concentrations ranging from 20, 40, 80, 120, 160, and 200 mg/L, respectively. As for the control, a native expression was set up such that Met with a final concentration of 100 mg/L was added on Day 2. Cells were 21 harvested on Day 5. The recombinant Mediator Head module was affinity-purified ^21^ and the protein complex yield at each condition was quantified. Plot of SeMet incorporation rate (%) into recombinant Mediator Head module expressed in Hi5 cells (A) or Sf9 cells (C) as a function of the concentration of SeMet added to the growth medium. Protein complex yield of the Mediator Head module expressed in Hi5 cells (B) or Sf9 cells (D) at each SeMet (or Met) concentration condition was plotted. Native: native expression where Met was added to a final concentration of 100 mg/L on Day 2; final concentrations of SeMet added on Day 2 is indicated in the plot.

Next, we conducted the same experiment in order to find out the optimal SeMet concentration for expression of the Mediator Head module in Sf9 cells. Since Sf9 cells appear to accommodate more SeMet than Hi5 cells (Figure 2) ^7^, we started the SeMet titration from 160 mg/L all the way up to 280 mg/L (Figure 5C, D). In this range, 160-280 mg/L, SeMet incorporation reached 75% at 200 mg/L and there is no increase at higher concentration, suggesting that 75% is the upper limit for SeMet incorporation, consistent with the previous report ^7^. The recovery of SeMet labeled Mediator Head module decreased as the concentration of SeMet increases (Figure 5D). Considering SeMet incorporation rate as well as the complex recovery rate, we concluded that for Sf9 cells, the optimal concentration of SeMet is 200 mg/L without overly compromising the recovery of the protein complex and achieving 75% SeMet incorporation.

### Application of our SeMet labeling method to multi-protein complexes involved in RNA polymerase II transcription

We further tested a generality of our labeling method by applying it to three other 22,23 multi-protein complexes involved in RNA polymerase II transcription ^22, 23^: human Taf8-Taf10 heterodimer with a molecular mass of 81kDa, yeast Mediator Middle module composed of seven subunits with a molecular mass of 190 kDa, and yeast TFIIF composed of three subunits with a molecular mass of 156 kDa, respectively. In our previous studies, the expression levels of human Taf8-Taf10, and yeast Mediator Middle module were fairly similar from low eMOI (0.5) to high eMOI (4.0) under native expression conditions. In fact, the optimal expression for the Mediator Middle module occurred with an eMOI= 0.5 ^21^. Yeast TFIIF was chosen to test the SeMet-labelling in Sf9 cells because it is one of only a few protein complexes we tested so far that expresses better in Sf9 cells than Hi5 cells ^21^. Following our protocol (supplemental protocol), Hi5 cells or Sf9 cells were cultured in Met-free medium for 24 hours followed by addition of the baculoviruses expressing each of these complexes with eMOIs ranging from 0.5 to 4.0. SeMet was added to each culture at a final concentration of 160 mg/L SeMet for Hi5 cells, and 200 mg/L for Sf9 cells 24 hours after infection. In parallel, native expressions using the same eMOI were set up as controls. The protein complexes were purified, yields of protein complexes were determined, and the extents of SeMet incorporation were measured (Figure 6).

**Figure 6:**
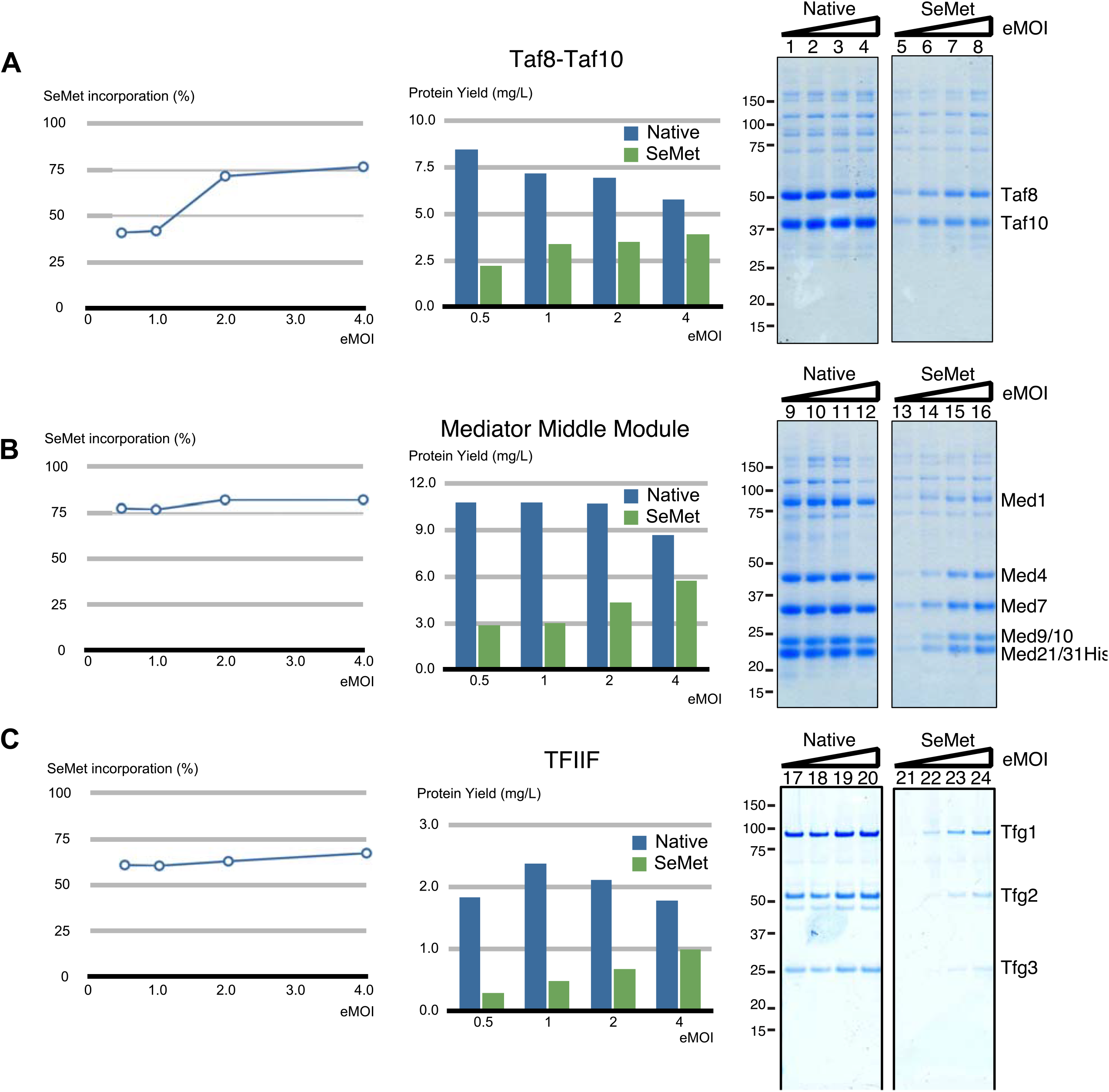
Application of the SeMet labeling method to human Taf8-Taf10, yeast Mediator Middle module, and yeast TFIIF complexes involved in RNA polymerase II transcription. SeMet-labeling method, “SeM-TEQC”, used for the Mediator Head module was applied to human Taf8-Taf10, the yeast Mediator Middle module, and yeast TFIIF. Human Taf8-Taf10, and the yeast Mediator Middle module were expressed in Hi5 cells, and yeast TFIIF was expressed in Sf9 cells. Following the experimental scheme described in Figure 3A, Hi5 or Sf9 cells (200 ml in 1-L shake flask) underwent Met depletion process. On Day 1, the recombinant baculovirus expressing human Taf8-Taf10, the yeast Mediator Middle module, or yeast TFIIF at an eMOI ranging from 0.5, 1.0, 2.0 and 4.0 was added followed by addition of Met at a final concentration of 100 mg/L as native expression control, or SeMet at a final concentration of 160 mg/L for Hi5 cells and 200 mg/L for Sf9 cells on Day 2. Cells were harvested on Day 5. For each condition, the three protein complexes were purified, the protein complex yields, as well as SeMet incorporation rates (%) were measured. Plot of SeMet incorporation rate (%) as a function of eMOI is displayed on the left; protein complex yields in native or SeMet-labeling expression as a function of eMOI is displayed in the middle; and SDS-PAGE of purified protein complex obtained from native or SeMet-labeling expression are on the right. Lanes 1-4: eMOI=0.5, 1.0, 2.0, 4.0; Lanes 5-8: eMOI=0.5, 1.0, 2.0, 4.0. (A) Human Taf8-Taf10, (B) yeast Mediator Middle module, and (C) yeast TFIIF

Consistent with our previous observation, the protein complex yields are fairly similar across different eMOI values for all three native complexes ^21^. Interestingly, the optimal eMOIs for all three complexes are at lower levels compared to the Mediator Head Module (Taf8-Taf10: 0.5; Mediator Middle module: 0.5; TFIIF: 1.0). However, in the presence of SeMet the complex yields expressed at a low eMOI were substantially reduced and the highest recovery was achieved with an eMOI of 4.0 (> 98% infectivity), which is in line with our observations for the Mediator Head module. Compared to the native expression, yields of the complexes are ranging from 56% for TFIIF to almost 70% for Taf8-Taf10 and Mediator Middle module with an eMOI of 4.0 and SeMet incorporation of these three complexes reached 68% to 78%. Taken together, these results clearly indicate that our labeling method is applicable to multiple systems. Since implementation of our TEQC protocol is key for this SeMet-labeling method, we name it SeM-TEQC method.

### Validation of our 24h-increment labeling protocol

One element of our experimental set up is a 24-hour infection period for the baculovirus prior to the addition of SeMet. If the Mediator Head module is expressed in this period, it would result in a production of non-SeMet-labeled protein complex, which may contribute to lowering the overall SeMet incorporation rate and may explain the maximal labeling of 75% we observed. Cronin et al. ^7^ indicated that addition of SeMet within the first 16 hours following viral infection is critical for reducing a production of non-SeMet-labeled protein - a key point in their report. Therefore, we decided to examine how much of the protein complex is expressed in the first 24 hours and thereafter. Following the experiment scheme illustrated in Figure 3A, we monitored expression of the Mediator Head module throughout the incubation period (Day 1-5) by western blotting using antibodies against 10His-Med17 (Δ1-108), Med18 (Δ109-140), and Med11 subunits of the Mediator Head module. In this experiment, every 24 hours, a small aliquot from the culture was taken at 24 hours intervals, and expressions of the three representative subunits mentioned above were probed by western blotting. As for a control, we set up a native expression of the Mediator Head module to which Met was added on Day 2 instead of SeMet. In the first 24 hours after addition of the virus, there was no detectable expression of any one of the three probed subunits in either the SeMet or native expression cultures (Figure 7). These data argue against the idea of a substantial production of non-SeMet-labeled protein complex between Day 1 and 2, further substantiating our 24h-increment labeling protocol.

**Figure 7:**
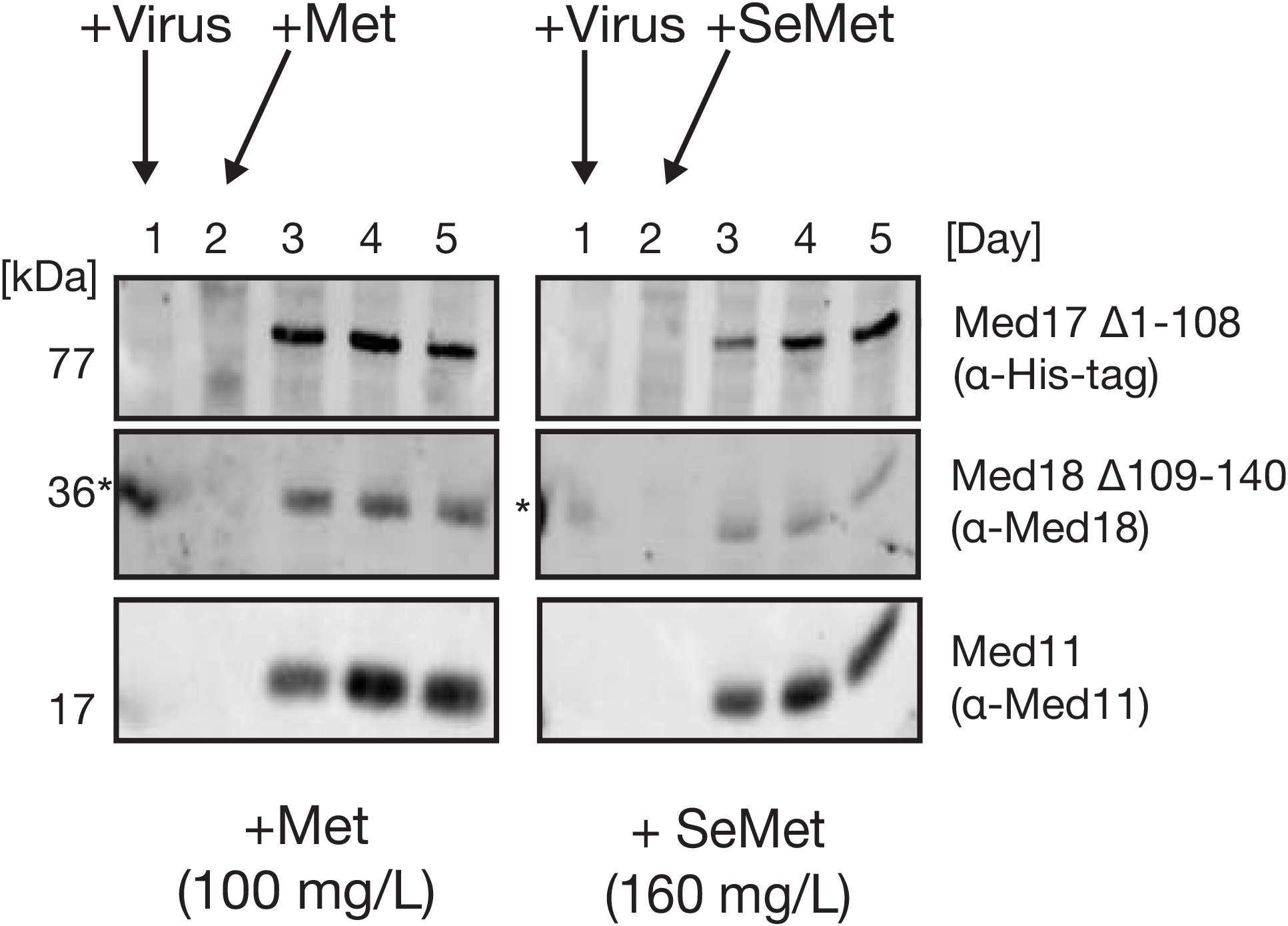
Expression profile of the Mediator Head Module during SeMet-labeling. Mediator Head Module production over the course of a standard SeMet labeling procedure (Figure 3A) compared to native expression was monitored by Western Blot analysis. Hi5 cells were grown in a Met-free medium for 24 hours, followed by virus infection (Day 1). After a 24-hour incubation period, either Met or SeMet was added to a final concentration of 100 mg/L or 160 mg/L, respectively, and the cells cultured for another 72 hours. Cell samples were collected every 24 hours (Day 1 to Day 5), lysed and subjected for SDS-PAGE followed by Western blotting. Western Blot of three representative subunits, Med17, Med18 and Med11, of the Mediator Head Module probed with either anti-His (Med17), anti-Med18 and anti-Med11, respectively. Time points of virus as well as Met or SeMet addition are indicated by arrows. Molecular mass (kDa) of each subunit is indicated on the left. (*) Signal bleed-over from adjacent marker lane

### Use of SeMet-labeled Mediator Head module for X-ray crystallography

The SeMet-labeled Mediator Head module was crystallized under similar conditions as the native Mediator Head module. The SeMet-labeled crystals were isomophorous with native crystals with a similar size. SeMet-containing crystals diffracted to 4.3 Å, and so did a representative native crystal. A scan of X-ray-induced fluorescence on the SeMet crystals demonstrated an absorption peak at 0.97948 A° (12,662 eV), consistent with the presence of SeMet. Diffraction data were collected at this wavelength, and processed with HKL2000 ^24^ as described previously ^4^. The initial phases were determined by SIRAS using the Ta_6_Br_14_ derivative or SAD by using the K_3_Ir(NO_3_)_6_ derivative. The phasing results were quite similar for both derivatives. The initial phases were extended by density modification by program PARROT, using SeMet datasets descried previously ^4^. On the last step of the phasing, 98 SeMet peaks of a possible 141 sites (~70%) in the unit cell were identified manually, and were used to calculate the experimental phases by the program PHASER ^25^. The final experimentally phased map and the SeMet peaks were used for model building of the Mediator Head module as described previously ^4^

To evaluate anomalous differential Fourier map peak positions with SeMet positions, peaks were researched by CCP4 program FFT from anomalous difference Fourier peak height from more than 3.5σ to 8σ, and summarized in Table 1. A total of 97 SeMet peaks were higher than 4.0σ in 20-6 A°, suggesting high quality of anomalous signals from Selenium in the crystals. It should be noted that CCP4 program 26 FFT ^26^ identified 97 SeMet peaks within a unit cell while 98 were identified manually.

**Table 1.** Peaks in 20-6 Å Anomalous Difference Fourier Map

|  | SeMet | Noise |
| --- | --- | --- |
| $\sigma$ Level | Peaks | Peaks |
| <8 | 44 | 0 |
| <7 | 64 | 0 |
| <6 | 76 | 0 |
| <5 | 89 | 0 |
| <4 | 97 | 10 |
| <3.5 | 97 | 77 |

To further validate Selenium peaks, we compared the known positions of methionine residues in the high-resolution structure of Med18-Med20, sub-complex of the Mediator Head module previously determined by X-ray analysis ^4^ with the Selenium anomalous peaks by placing Med18-Med20 structure onto the 4.3 Å Mediator Head module map. Selenium anomalous difference Fourier map peak heights of Med18 and Med20 were summarized in Table 2. Peaks ranging from 3.0 σ to 9.6 σ were observed for 17 of possible 27 methionine sites for Med18, and 5 of possible 9 methionine sites for Med20 were observed: 22 of a possible 36 sites (61%) (Table 2) -the methionine residues at N-termini of both subunits were not included. Anomalous difference Fourier map peaks calculated from 20-6 Å at contoured at 3 σ, nicely correlate with the location of known methionine sites in all 3 molecules found in the asymmetric unit (Figure 8A-C).

**Table 2.** Se Anomalous Difference Fourier Map Peak Hight of Med18

| Met residue Number | Mol1 | Mol2 | Mol3 |
| --- | --- | --- | --- |
| 1 | ND | ND | ND |
| 80 | ND | 7.0 | ND |
| 81 | 3.8 | 4.4 | 6.0 |
| 104 | ND | ND | ND |
| 150 | ND | ND | ND |
| 182 | 6.3 | 7.2 | 5.9 |
| 204 | 6.6 | 8.7 | ND |
| 224 | 6.6 | 8.7 | 6.9 |
| 276 | 4.9 | 4.7 | 3.9 |
| 296 | ND | 4.5 | 3.0 |

Se Anomalous Difference Fourier Map Peak Hight of Med20
| Met residue Number | Mol1 | Mol2 | Mol3 |
| --- | --- | --- | --- |
| 1 | ND | ND | ND |
| 57 | ND | 9.6 | 6.1 |
| 78 | ND | 6.8 | ND |
| 117 | ND | 4.8 | 3.5 |

**Figure 8:**
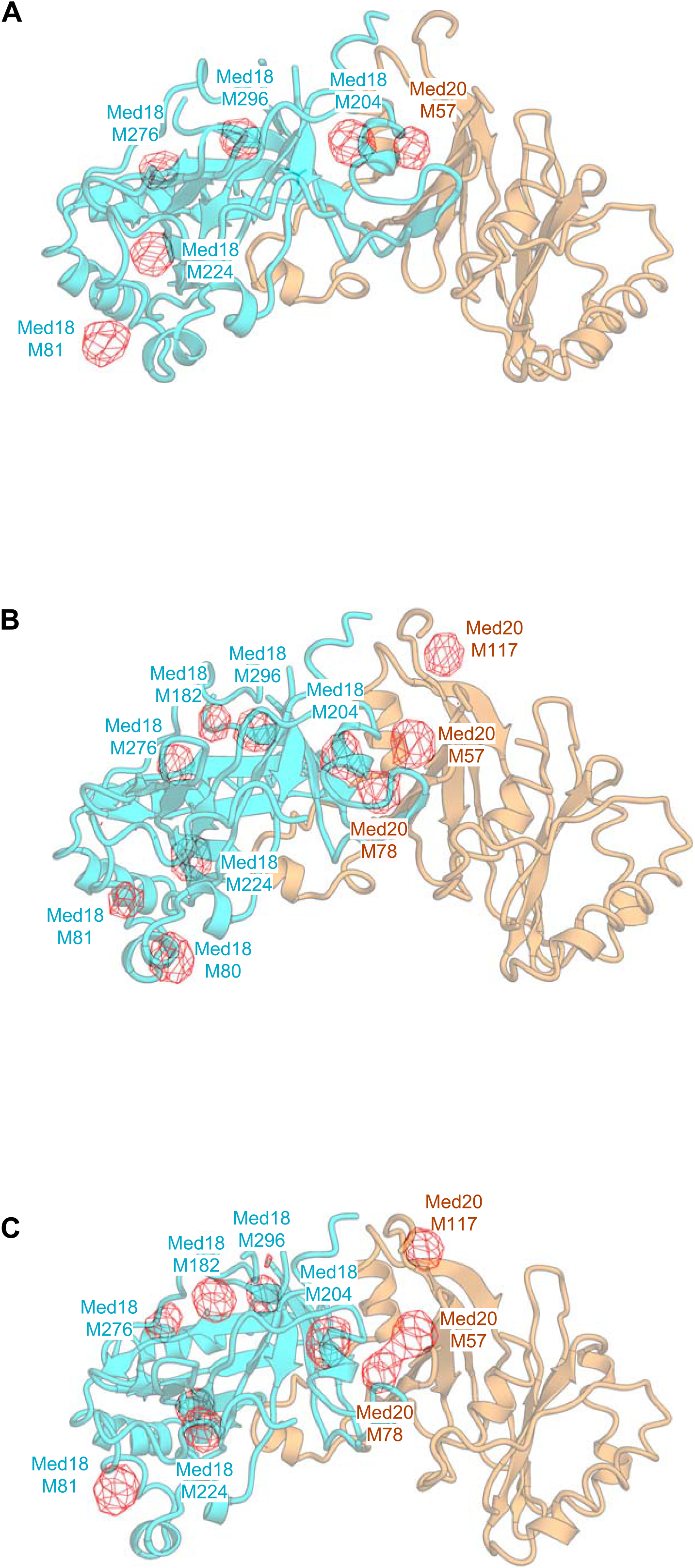
Comparison between Se anomalous peaks on the Med18-Med20 structure, and the known positions of methionine residues in the Med18-Med20 structure. Se Anomalous Difference Fourier Map was superimposed onto the cartoon view of the high-resolution structure of the Med18-Med20 structure determined in the previous study ^38^. The Selenium anomalous difference Fourier map calculated from 20-6 Å at contoured at 3 σ is shown in red and overlaid onto Med18-Med20. Three molecules in the asymmetric unit are displayed individually as (A)-(C). The height of the anomalous peaks is shown in Table 2. M: methionine residue (e.g. M57: methionine residue 57)

## Discussion

In this work, we developed a simple and robust protocol termed SeM-TEQC method for productions of SeMet-labeled protein complexes in the insect cells by addressing the issues related to (i) effect of Met depletion on the cells, (ii) timing of addition of SeMet, and most importantly, (iii) SeMet toxicity toward the insect cells. Our first finding was that the insect cells tolerate Met depletion for an extended period of time under otherwise normal growth conditions (Figure 1), making medium exchanges with dialyzed FBS, or any other supplemental reagents during methionine starvation as reported previously ^6,17^ unnecessary. Second, based on our analysis, no detectable protein expression was observed 24 hours following the virus infection (Figure 7), eliminating the need for SeMet addition 16 hours after viral infection ^7^ and permitting a more feasible 24-hour schedule. Finally, and most importantly, we addressed the issue of the SeMet toxicity in the insect cells. This work outlines cell line dependent toxicity thresholds and establishes that the inherent SeMet toxicity can be evaded by optimizing baculovirus infection levels. Our previously developed TEQC method proved as an essential tool to limit SeMet toxicity as such it allows us to quantitatively control virus infection levels in a convenient and easy manner, and is crucial for optimal SeMet incorporation. In brief, to achieve full baculovirus infection, the eMOI needs to be 4.0 or higher, which corresponds to ~98% infectivity ^21^. In retrospect, the condition used by Chen and Bahl corresponds to an MOI of 5-10 for SeMet-labeleling of human choriogonadotropin (hCG) in the insect cells, which turns out to be in the optimal range to evade SeMet toxicity, even though the authors failed to provide the reasoning to their MOI choice ^17^. Similarly, a high MOI was used in the works of Carfi et al. ^27^ and Bellizzi et al. ^6^ without any explanation as to why. On the contrary, Cronin et al. used a baculovirus MOI of 1 for infection, which is, according to our results, less optimal to evade SeMet toxicity and might explain the low recovery of SeMet-labeled proteins they encountered ^7^.

### Possible mechanism of SeMet toxicity

Mechanism of SeMet toxicity toward eukaryotic cells has been studied mostly using the yeast *Saccharomyces cerevisiae* ^10, 18^. As these studies indicated, SeMet toxicity appears to come from two different mechanisms: SeMet likely generates reactive species, resulting in DNA damage ^28^, consistent with the observation that SeMet could effectively inhibit the insect cell growth ^17^. Alternatively, SeMet toxicity results from its metabolites, and a random incorporation of which promotes protein aggregation - SeMet causes proteotoxic stress resulting in cell death ^29^. How does baculovirus infection reduce SeMet toxicity? As reported, baculovirus infection results in cell cycle and DNA replication arrest ^30,31^. Once infected, the insect cells are no longer able to divide. As a result, the effect of SeMet-induced DNA damage could be neutralized, thereby maintaining overall cell viability as we observed (Figures 3, 4).

Considering the mechanism of SeMet toxicity being attributed to SeMet metabolites, we reasoned that SeMet toxicity could be minimized by reducing the amount of toxic SeMet metabolites (e.g. Selenocysteine) through impairing conversion of Se-adenosylselenomethionine (SeAM) to Se-adenosylselenohomocysteine (SeAH) ^32^. In fact, this strategy worked well in the yeast S. *cerevisiae* by knocking out genes (SAM1, SAM2, or CYS3) encoding enzymes S-adenosylmethionine synthetase, or Cystathionine gamma-lyase in the pathway that produces toxic SeMet metabolites ^10,18^. Accordingly, we attempted to apply a similar idea to disrupt conversion of SeMet to SeAM by addition of SAM to the medium, which we hoped competes with SeAM as a substrate for methyltransferases, thereby minimizing the synthesis of SeMet metabolites. However, addition of SAM to the growth medium did not reduce overall SeMet toxicity (data not shown). Thus, supplementing SAM to the insect cell culture may not effectively disrupt the metabolic pathway that produces SeMet metabolites as 10,18 shown in yeast ^10,18^. Thus, the main SeMet toxic effect may result from SeMet-induced DNA damage and not via toxic SeMet metabolites. Alternately, it might be simply that a complete viral infection is forcing all excess SeMet toward incorporation into the recombinant protein, avoiding toxic cellular effects seen for uninfected cells.

Regardless, deciphering the exact mechanisms is beyond the scope of this work and will have to be addressed at a later time.

### Generality of the SeM-TEQC method

The Mediator Head module was used as a model protein complex to develop the SeM-TEQC method with subunits composition ranging from 15 kDa to 77 kDa. Moreover, we have shown applicability of our method by expanding our protocol to label human Taf8-Taf10 heterodimer with a molecular mass of 81 kDa, yeast Mediator Middle composed of seven subunits with a molecular mass of 190 kDa, and yeast TFIIF composed of three subunits with a molecular mass of 156 kDa. There was no significant difference among these complexes in terms of SeMet incorporation level, and protein yield compared to their corresponding native expressions, supporting the generality of our method. Having focused primarily on expression and SeMet incorporation of multiprotein complex, application of our method on single-subunits has been successfully achieved and led to the structure determination of the tight junction protein Claudin by SeMet SAD phasing ^33^, again proving the versatility of the SeM-TEQC method.

As Cronin et al. pointed out ^7^, the advantage of the existing methods for secreted proteins, including engineered secrete expression ^6,17^, lies in its ability to remove unlabeled proteins by medium exchange during the procedure, thereby increasing overall SeMet incorporation. For example, Chen and Bahl reported^17^ that SeMet incorporation rate of human choriogonadotropin (hCG) reached 84%, which is better than what our SeM-TEQC method (~75%) achieved. However, the report by Bellizzi et al.^6^ - this publication that has been widely cited for SeMet protocol for BEVS in the insect cells - indicated that their SeMet incorporation rate was 76%, which is comparable to that of the SeM-TEQC method. However, our protocol was developed primarily for intracelluar multi-protein complexes. It has not been applied to secreted proteins yet. It still remains to be seen how well SeM-TEQC method works for those. Based on our current results, we would expect it to work equally good if not better than the existing methods

## Conclusions

Our discovery that SeMet toxicity can be circumvented by a high baculoviral infection led us to develop a simple and quantitative SeMet-labeling method termed SeM-TEQC. The overall scheme is illustrated in Figure 9. This method does not require laborious procedures (medium exchange during the procedure) or additional reagents (e.g. dialyzed FBS). Thus, this method is more cost effective compared to the exiting protocols. The protocol steps are carried out in 24 hours increments, making the procedure practical and easy to implement. Our SeM-TEQC method enables an optimal production of SeMet-labeled proteins or protein complexes with SeMet incorporation levels to about 75% and protein yield of 60-90% compared to the native protein expression.

**Figure 9:**
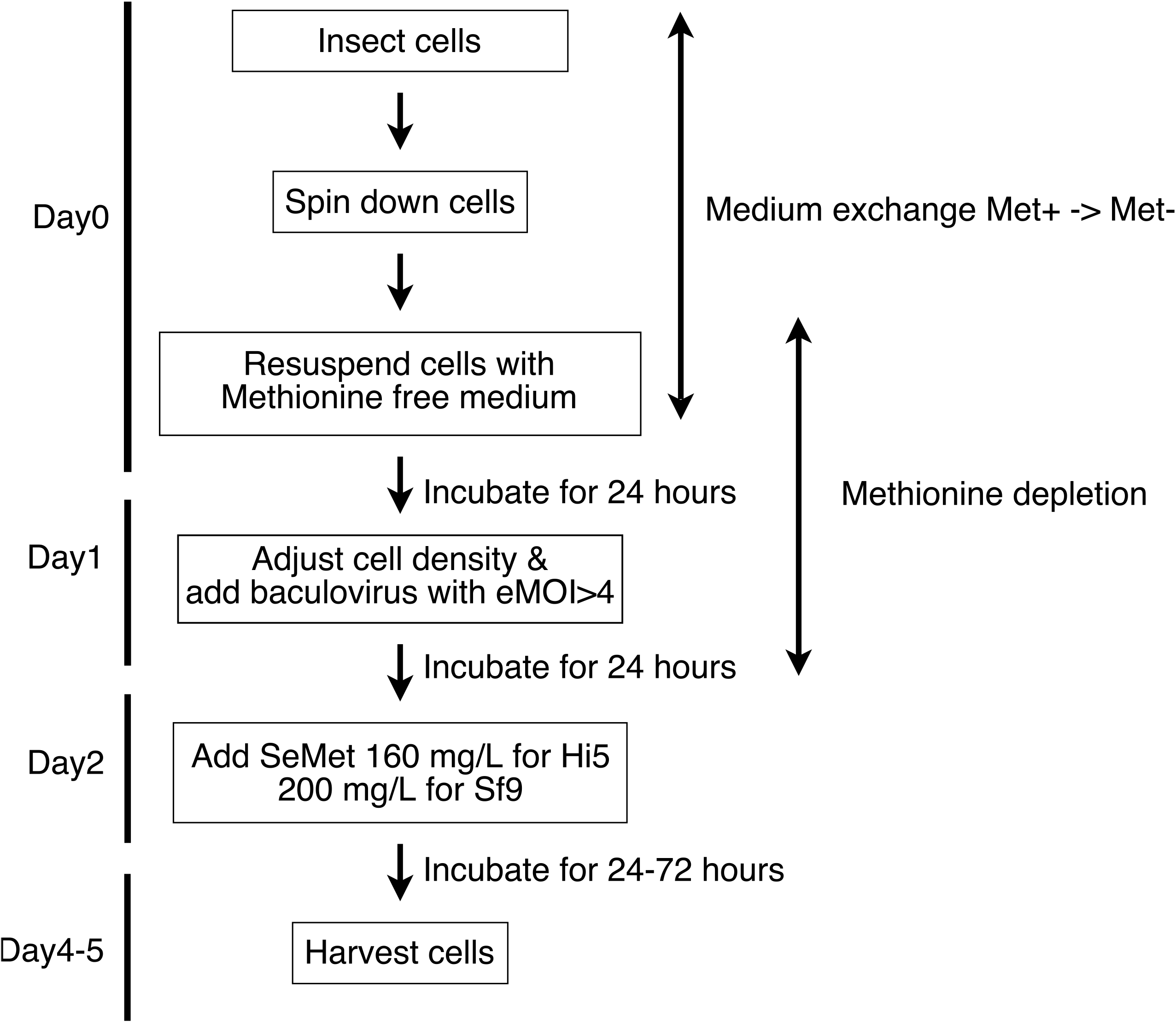
The overall experimental scheme for SeM-TEQC method.

## Materials and Methods

### Maintenance of the insect cells, recombinant baculovirus production, virus storage, and virus titer estimation

Cells are maintained as described in ^21^. Generation of recombinant baculoviruses was described in ^21,34^. Generation of frozen virus stocks was described in ^21,35^. The titer estimates of the recombinant baculoviruses (eTiters) were determined by TEQC method as described in ^21^.

### Cell proliferation assay for methionine depleted conditions

Healthy dividing Hi5 or Sf9 insect cells were centrifuged in a 50 ml conical tube and methionine-containing medium (ESF921) (Expression systems, Davis CA) was discarded. Cells were resuspended in Met-free ESF921 medium (delta series, methionine deficient) (Expression systems, Davis CA), or as a control, methionine containing medium (ESF921) to set up a 50 ml culture with a cell density of 0.5 x 10^6^ cells/ml for Hi5 cells or 0.6 x 10^6^ cells/ml for Sf9 cells. Cell cultures were incubated on a shaker at 125 rpm at 27 °C. Every 24 hours for up to 96 hours, the cell density as well as cell viability were measured with a TC20™ Automated Cell counter (Bio-Rad).

### Cell proliferation assay in presence of SeMet

Healthy dividing Hi5 or Sf9 insect cells were centrifuged in a 50 ml conical tube and methionine-containing medium (ESF921) was discarded. Cells were resuspended in methionine-free medium (ESF921), and as a control, methionine-containing medium to set up a total of 7 flasks of 50 ml cultures with the cell density of 0.5 x 10^6^ cells/ml for Hi5 cells or 0.6 x 10^6^ cells/ml for Sf9 cells. Sets of cultures in methionine-containing medium or in methionine-depleted medium were set up in parallel. SeMet was added to each culture at the concentration indicated in the figure legends. Cell cultures were incubated at 27 °C while shaking at 125 rpm. Every 24 hours, the cell density and viability were measured with a TC20™ Automated Cell counter (Bio-Rad).

### Expressions of native and SeMet-labeled protein complexes

Expressions of native multi-protein complexes were performed in 200 ml cultures with 1.0 x 10^6^ cells/ml of Hi5 cells or 1.5 x 10^6^ cells/ml of Sf9. Cultures were infected with the recombinant baculoviruses at the indicated eMOI, which was calculated by our TEQC method ^21^. Each culture was incubated at 27 °C for 96 hours. The expression of SeMet-labeled protein complexes is described in detail in the Supplemental protocol. Briefly, for medium exchange, Hi5 or Sf9 insect cells were centrifuged, methionine containing medium was discarded, and cells resuspended in 200 ml methionine-depleted medium ESF921 to a cell density of 1.0 x 10^6^ cells/ml for Hi5 cells or 1.5 x 10^6^ cells/ml for Sf9 cells. Cell cultures were incubated on a shaker at 125 rpm for 24 hours at 27 °C in order to deplete endogenous methionine. After 24 hours, cell density was adjusted to 1.0 x 10^6^ cells/ml in 200 ml for Hi5 cells or 1.5 x 10^6^ cells/ml for Sf9 with methionine-free medium. At this point, cells were infected with baculovirus at the various eMOI values indicated in the main text. After incubating for another 24 hours at 27 °C, SeMet was added to a final concentration of 20-200 mg/L and cells cultured for 72 hours. Cells were harvested by centrifugation, cell pellets were frozen in liquid nitrogen, and stored at -80 °C until use.

### Purification of protein complexes

Purification procedures for the Mediator Head module, Taf8-Taf10, TFIIF, and Mediator middle module were described previously ^21^.

### Determination of SeMet incorporation by amino acid analysis (AAA)

The incorporation of SeMet was determined by amino acid analysis carried out at The University of California Davis proteomics core facility. The details of AAA are described in (http://msf.ucdavis.edu/amino-acid-analysis/). Briefly, since Met is destroyed during hydrolysis with 6 N HCl, Met was determined by converting it to the acid stable form, methionine sulfone, with oxidation using performic acid, prior to the standard acid hydrolysis ^36^. Quantity of each amino acid was measured by AAA. Since valine (Val), leucine (Leu), and phenylalanine (Phe) are least affected by the oxidation process with performic acid, we used the average quantity of these three amino acids to compare the Met ratio in native and SeMet-labeled proteins. The percentage of SeMet incorporation was calculated using the following equation:

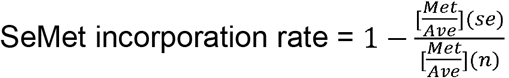, where Met (n): quantity of Met under a native ö«) expression; Ave (n): average quantity of Val, Leu, and Phe combined under a native expression; Met (se): quantity of Met under SeMet-labeling expression; Ave (se): average quantity of Val, Leu, and Phe combined under SeMet-labeling expression. The detailed description of how this equation was derived is described in Supplemental Materials. The ratio in quantity between Met and the average value of Val, Leu, and Phe under native as well as SeMet-labeling expression condition was first calculated followed by determining the incorporation rate using the equation described above.

### X-ray crystallography for the SeMet-labeled Mediator Head module

The SeMet-labeled Mediator Head module crystals were obtained by the hanging-drop vapor-diffusion method as described previously ^4^. Diffraction data were collected at beamline 23ID at the Advanced Photon Source (APS) at Argonne National Laboratory. All diffraction data were processed with HKL2000. The structure was determined by selenomethionine (SeMet) single-wavelength anomalous dispersion (SAD) after a sufficient number (98) of SeMet sites had been identified by a combination of the initial phases from Ta_6_Br_14_ and Iridium derivatives, and partial model SAD phases as described previously ^4^.

### Western blotting

Hi5 cells were infected with the recombinant baculovirus expressing the Mediator Head module with eMOIs = 4.0 after Met depletion. 24 hours after addition of the virus, SeMet was added to the culture at a final concentration of 160 mg/L, and incubated for an additional 3 days. 1 ml aliquot of cell culture was taken every 24 hours, and cells were harvested in 1.5 ml tubes. Cell pellets were frozen in liquid nitrogen, and stored at -80 °C until use. Preparation of cell lysate and the method for western blotting is described in ^21^. The blot was probed for the Head module with anti-His tag monoclonal mouse antibody (Thermo Scientific Pierce) for 10xHis-tagged Med17, as well as with rabbit anti-Med18 (anti-Srb5) and rabbit anti-Med11 antibodies ^37^. Detection was carried out with a ChemiDoc MP imaging system (BioRad) using Dylight 680 goat anti-rabbit IgG (Thermo Scientific Pierce) for Med18 and Med11, and Dylight 800 goat anti-mouse IgG (Thermo Scientific Pierce) for anti-His tag as secondary antibodies.

## Supplemental Material

Calculation of the SeMet incorporation rate using AAA data, and supplemental protocol for the SeM-TEQC method

## Acknowledgments

We thank Dr. T. Hurley for critical reading of the manuscript. We thank Dr. John Schulze at The University of California Davis proteomics core facility for conducting amino acid analysis, and for providing the technical information. This research was supported by US National Science Foundation grant MCB-1157688, the National Institutes of Health (R01 GM111695), and Showalter Trust Fund to (Y.T.), Japan Science and Technology Agency (PRESTO, JPMJPR14L2) to (T.I.). X-ray data were collected at the GM/CA-CAT at the Advanced Photon Source (APS), Argonne National Laboratory, Argonne, IL. GM/CA-CAT is funded by National Cancer Institute grant Y1-CO-1020 and National Institute of General Medical Sciences grant Y1-GM-1104. Use of the Advanced Photon Source was supported by the U.S. Department of Energy, Basic Energy Sciences, Office of Science, under contract DE-AC02-06CH11357.

## Abbreviations and symbols

AAA: Amino Acid Analysis
BEVS: baculovirus expression vector system
*E. coli*: *Escherichia coli*
FBS: Fetal bovine serum
hCG: human choriogonadotropin
Hi5: *Trichoplusia ni*
Leu: L-leucine
Met: L-methionine
Met+: L-methionine-containing
Met-: L-methionine-free
MAD: multi-wavelength anomalous diffraction
MOI: multiplicity of infection
eMOI: estimated multiplicity of infection
Phe: L-phenylalanine
SAD: single-wavelength anomalous diffraction
Sf9: *Spodoptera frugiperda*
Se: Selenium
SeMet: Seleno-L-methionine
SeAM: Se-adenosylselenomethionine
SeAH: Se-adenosylselenohomocysteine
SeCys: Selenocysteine
S. *cerevisiae*: *Saccharomyces cerevisiae*
TEQC method: Titer Estimation for Quality Control method
Val: L-valine

## References

1. Hendrickson WA, Horton JR, LeMaster DM (1990) Selenomethionyl proteins produced for analysis by multiwavelength anomalous diffraction (MAD): a vehicle for direct determination of three-dimensional structure. EMBO J 9:1665–1672.

2. Hendrickson WA, Ogata CM (1997) [28] Phase determination from multiwavelength anomalous diffraction measurements. Methods Enzymol 276:494–523.

3. Bushnell DA, Cramer P, Kornberg RD (2001) Selenomethionine incorporation in Saccharomyces cerevisiae RNA polymerase II. Structure 9:R11–14.

4. Imasaki T, Calero G, Cai G, Tsai KL, Yamada K, Cardelli F, Erdjument-Bromage H, Tempst P, Berger I, Kornberg GL, Asturias FJ, Kornberg RD, Takagi Y (2011) Architecture of the Mediator head module. Nature 475:240–243.

5. Kitajima T, Yagi E, Kubota T, Chiba Y, Nishikawa S, Jigami Y (2009) Use of novel selenomethionine-resistant yeast to produce selenomethionyl protein suitable for structural analysis. FEMS Yeast Res 9:439–445.

6. Bellizzi JJ, Widom J, Kemp CW, Clardy J (1999) Producing selenomethionine-labeled proteins with a baculovirus expression vector system. Structure 7:R263–267.

7. Cronin CN, Lim KB, Rogers J (2007) Production of selenomethionyl-derivatized proteins in baculovirus-infected insect cells. Protein Sci 16:2023–2029.

8. Barton WA, Tzvetkova-Robev D, Erdjument-Bromage H, Tempst P, Nikolov DB (2006) Highly efficient selenomethionine labeling of recombinant proteins produced in mammalian cells. Protein Sci 15:2008–2013.

9. Carfi A, Gong H, Lou H, Willis SH, Cohen GH, Eisenberg RJ, Wiley DC (2002) Crystallization and preliminary diffraction studies of the ectodomain of the envelope glycoprotein D from herpes simplex virus 1 alone and in complex with the ectodomain of the human receptor HveA. Acta Crystallogr D Biol Crystallogr 58:836–838.

10. Malkowski MG, Quartley E, Friedman AE, Babulski J, Kon Y, Wolfley J, Said M, Luft JR, Phizicky EM, DeTitta GT, Grayhack EJ (2007) Blocking S-adenosylmethionine synthesis in yeast allows selenomethionine incorporation and multiwavelength anomalous dispersion phasing. Proc Natl Acad Sci U S A 104:6678–6683.

11. Kost TA, Condreay JP, Jarvis DL (2005) Baculovirus as versatile vectors for protein expression in insect and mammalian cells. Nat Biotechnol 23:567–575.

12. Massotte D (2003) G protein-coupled receptor overexpression with the baculovirus-insect cell system: a tool for structural and functional studies. Biochim Biophys Acta 1610:77–89.

13. Assenberg R, Wan PT, Geisse S, Mayr LM (2013) Advances in recombinant protein expression for use in pharmaceutical research. Curr Opin Struct Biol 23:393–402.

14. McWhirter SM, Pullen SS, Holton JM, Crute JJ, Kehry MR, Alber T (1999) Crystallographic analysis of CD40 recognition and signaling by human TRAF2. Proc Natl Acad Sci U S A 96:8408–8413.

15. Kornberg RD (2005) Mediator and the mechanism of transcriptional activation. Trends Biochem Sci 30:235–239.

16. Cai G, Imasaki T, Yamada K, Cardelli F, Takagi Y, Asturias FJ (2010) Mediator head module structure and functional interactions. Nature structural & molecular biology 17:273–279.

17. Chen WY, Bahl OP (1991) Selenomethionyl analog of recombinant human choriogonadotropin. J Biol Chem 266:9355–9358.

18. Bockhorn J, Balar B, He D, Seitomer E, Copeland PR, Kinzy TG (2008) Genome-wide screen of Saccharomyces cerevisiae null allele strains identifies genes involved in selenomethionine resistance. Proc Natl Acad Sci U S A 105:17682–17687.

19. Fremont DH, Crawford F, Marrack P, Hendrickson WA, Kappler J (1998) Crystal structure of mouse H2-M. Immunity 9:385–393.

20. Liemann S, Chandran K, Baker TS, Nibert ML, Harrison SC (2002) Structure of the reovirus membrane-penetration protein, Mu1, in a complex with is protector protein, Sigma3. Cell 108:283–295.

21. Imasaki T, Wenzel S, Yamada K, Bryant ML, Takagi Y (2018) Titer estimation for quality control (TEQC) method: A practical approach for optimal production of protein complexes using the baculovirus expression vector system. PLoS One 13:e0195356.

22. Boeger H, Bushnell DA, Davis R, Griesenbeck J, Lorch Y, Strattan JS, Westover KD, Kornberg RD (2005) Structural basis of eukaryotic gene transcription. FEBS Lett 579:899–903.

23. Galbraith MD, Donner AJ, Espinosa JM (2010) CDK8: a positive regulator of transcription. Transcription 1:4–12.

24. Otwinowski Z, Minor W (1997) Processing of X-ray Diffraction Data Collected in Oscillation Mode. Methods in Enzymology 276:307–326.

25. McCoy A, Grosse-Kunstleve R, Adams P, Winn M, Storoni L, Read R (2007) Phaser crystallographic software. J Appl Crystallogr 40:658–674.

26. Read RJ, Schierbeek AJ (1988) A phased translation function. Journal of Applied Crystallography 21:490–495.

27. Carfi A, Willis SH, Whitbeck JC, Krummenacher C, Cohen GH, Eisenberg RJ, Wiley DC (2001) Herpes simplex virus glycoprotein D bound to the human receptor HveA. Mol Cell 8:169–179.

28. Seitomer E, Balar B, He D, Copeland PR, Kinzy TG (2008) Analysis of Saccharomyces cerevisiae null allele strains identifies a larger role for DNA damage versus oxidative stress pathways in growth inhibition by selenium. Mol Nutr Food Res 52:1305–1315.

29. Plateau P, Saveanu C, Lestini R, Dauplais M, Decourty L, Jacquier A, Blanquet S, Lazard M (2017) Exposure to selenomethionine causes selenocysteine misincorporation and protein aggregation in Saccharomyces cerevisiae. Scientific reports 7:44761.

30. Braunagel SC, Parr R, Belyavskyi M, Summers MD (1998) Autographa californica nucleopolyhedrovirus infection results in Sf9 cell cycle arrest at G2/M phase. Virology 244:195–211.

31. Ikeda M, Kobayashi M (1999) Cell-cycle perturbation in Sf9 cells infected with Autographa californica nucleopolyhedrovirus. Virology 258:176–188.

32. Lazard M, Dauplais M, Blanquet S, Plateau P (2015) Trans-sulfuration Pathway Seleno-amino Acids Are Mediators of Selenomethionine Toxicity in Saccharomyces cerevisiae. J Biol Chem 290:10741–10750.

33. Suzuki H, Nishizawa T, Tani K, Yamazaki Y, Tamura A, Ishitani R, Dohmae N, Tsukita S, Nureki O, Fujiyoshi Y (2014) Crystal structure of a claudin provides insight into the architecture of tight junctions. Science 344:304–307.

34. Fitzgerald DJ, Berger P, Schaffitzel C, Yamada K, Richmond TJ, Berger I (2006) Protein complex expression by using multigene baculoviral vectors. Nat Methods 3:1021–1032.

35. Wasilko DJ, Lee SE, Stutzman-Engwall KJ, Reitz BA, Emmons TL, Mathis KJ, Bienkowski MJ, Tomasselli AG, Fischer HD (2009) The titerless infected-cells preservation and scale-up (TIPS) method for large-scale production of NO-sensitive human soluble guanylate cyclase (sGC) from insect cells infected with recombinant baculovirus. Protein Expr Purif 65:122–132.

36. Hirs CHW. Determination of cystine as cysteic acid. (1967) Methods in Enzymology. Academic Press, pp. 59–62.

37. Takagi Y, Kornberg RD (2006) Mediator as a general transcription factor. J Biol Chem 281:80–89.

38. Lariviere L, Geiger S, Hoeppner S, Rother S, Strasser K, Cramer P (2006) Structure and TBP binding of the Mediator head subcomplex Med8-Med18-Med20. Nat Struct Mol Biol 13:895–901.

