## Supplemental Materials for "A practical method for efficient and optimal production of selenomethionine-labeled recombinant protein complexes in the insect cells"

**Calculation of the SeMet incorporation rate using AAA data:**

SeMet incorporation rate can be described as:

SeMet incorporation rate = $\frac{SeMet(se)}{Met\left( se \right)+SeMet\left( se \right)}$ - Equation 1

where Met(se): quantity of Met, SeMet(se): quantity of SeMet under a SeMet-labeling expression condition

We assume that a ratio between a total Met, and Val, Leu, and Phe should stay the same under native or SeMet-labeling condition such that

$\frac{Met(n)}{Ave (n)}=\frac{Met\left( se \right)+SeMet(se)}{Ave (se)}=\frac{Met(se)}{Ave \left( se \right)}+\frac{SeMet(se)}{Ave(se)}$ - Equation 2

where Met (n): quantity of Met in native expression; Ave (n): average quantity of Val, Leu, and Phe combined under native expression condition; Ave (se): average quantity of Val, Leu, and Phe combined under SeMet-labeling expression condition.

Equation 2 can be modified such that

$\frac{SeMet(se)}{Ave (se)}=\frac{Met(n)}{Ave \left( n \right)}-\frac{Met(se)}{Ave(se)}$ - Equation 3

Equation 1 can be written as:

Incorporation rate = $\frac{SeMet(se)}{Met\left( se \right)+SeMet\left( se \right)}=\frac{\frac{SeMet(se)}{Ave(se)}}{\frac{Met(se)}{Ave(se)}+\frac{SeMet(se)}{Ave (se)}}$ - Equation 4

By combining Equations 3 and 4, SeMet incorporation rate can be described as:

SeMet incorporation rate = $1-\frac{[\frac{Met}{Ave}](se)}{[\frac{Met}{Ave}](n)}$ - Equation 5

The ratio in quantity between Met and the average value of Val, Leu, and Phe combined in the native as well as SeMet-labeling expression was first calculated followed by determining the incorporation rate using Equation 5.

**Supplemental Protocol for the SeM-TEQC Method:**

The detailed protocols for maintenance of cells, generation of recombinant baculoviruses, and generation of frozen virus stocks were described previously (2). Our SeM-TEQC method consists of 3 steps: (i) titer estimation of the recombinant baculovirus, (ii) depletion of endogenous methionine in the insect cells, (iii) SeMet labeling of the protein or protein complexes of interest. The protocol is as follows:

**(I) Titer estimation of the recombinant baculoviruses (eTiters, eMOI)**

The first step is to estimate the titer of the recombinant baculovirus expressing protein or protein complex of interest. This can be achieved using our TEQC method (2).

**(II) Procedure for depletion of endogenous methionine in the insect cells:**

The second step is to prepare the insect cells in which the endogenous methionine pool is sufficiently depleted. The protocol for setting up a 200 ml culture is described below. Expression can be scaled up.

**Day 0:**

1. Seed a 200 ml culture of healthy insect cells (Hi5, Sf9) in a 1 L ventilated flask to a density of 4-7 x 10^6^ cells/ml in Met-containing media (ESF921; Expression systems).
2. Transfer 2.0 x 10^8^ cells for Hi5 (corresponds to 1.0 x 10^6^ cells/ml in 200 ml), or 3.0 x 10^8^ cells for Sf9 (corresponds to 1.5 x 10^6^ cells/ml in 200 ml) in to 50 ml conical tubes, centrifuge for 5 min at 1,500 rpm at RT using Centra CL2 centrifuge (Thermo Fishers), and aspirate supernatant. For example, if the cell density of Hi5 cells is 5.0 x 10^6^ cells/ml, then 40 ml of this culture will provide the cells, which give rise to 200 ml culture with 1.0 x 10^6^ cells/ml.
3. Prepare a 1 L ventilated flask with 170 ml of methionine-free ESF921 media (ESF921 delta series, Methionine deficient, Expression Systems, Inc.)
4. Resuspend cells gently in about 30 ml of methionine-free ESF921 medium (ESF921 delta series, methionine deficient, Expression Systems, Davis, CA) and add cells to the prepared 1 L flask, mix gently. The final cell density is 1.0 x 10^6^ cells/ml for Hi5 cells, or 1.5 x 10^6^ cells/ml for Sf9 cells, respectively.
5. Incubate the cultures on shaker with 125 rpm, at 27 °C for 24 hours in order to deplete endogenous methionine. Cells will grow at the same rate as in a regular methionine-containing medium. After 24 hours, the cell density will usually be doubled.

**Day 1:**

6. Adjust cell density to 1.0 x 10^6^ cells/ml for Hi5 cells with methionine-free medium in a 1 L flask. As for Sf9 cells, adjust cell density to 1.5 x 10^6^ cells/ml accordingly. Add the recombinant baculovirus with an eMOI=4.0 or greater to ensure that the cells will be fully infected.

**(III) SeMet-labeling:**

The third and final step is to label expressed proteins with SeMet as follows:

**Day 2:**

7. Prepare 25mg/ml L-SeMet stock (purity > 98%, CHEM-IMPEX INT’L INC.) and filter sterilize.
8. Add L-SeMet to the culture at a final concentration of 160 mg/L for Hi5 cells, or 200 mg/L for Sf9 cells. Note that at this point, the insect cells have been deprived of methionine for 48 hours, and the cells should be fully infected.

**Day 4-5 and thereafter:**

9. Incubate culture on shaker at 100 rpm, 27 °C for 96 hours for viruses generated by the MultiBac system (1), or 72 hours for viruses generated with Bac to Bac or a similar system.

10. Purify the expressed protein or protein complexes for crystallization.

Note that MultiBac system utilizes the modified baculovirus genome in which the protease gene, *V-CATH*, important for the lytic cycle of virus, has been deleted, enabling a prolonged period for protein expression. That is why we harvest cells 96 hours instead of 72 hours after infection.

**References:**

1. Berger I, Fitzgerald DJ, Richmond TJ (2004) Baculovirus expression system for heterologous multiprotein complexes. Nat Biotechnol 22:1583-1587. PMID: 15568020 {Medline}

2. Imasaki T, Wenzel S, Yamada K, Bryant ML, Takagi Y (2018) Titer estimation for quality control (TEQC) method: A practical approach for optimal production of protein complexes using the baculovirus expression vector system. PLoS One 13:e0195356. PMID: 29614134 {Medline}
